# Characterization of a nosocomial outbreak caused by VIM-1 *Klebsiella michiganensis* using Fourier-Transform Infrared (FT-IR) Spectroscopy

**DOI:** 10.1101/2024.07.18.604080

**Authors:** David Rodriguez-Temporal, María Sánchez-Cueto, Sergio Buenestado-Serrano, Mario Blázquez-Sánchez, Emilia Cercenado, Mark Gutiérrez-Pareja, Andrea Molero-Salinas, Elena López-Camacho, Patricia Muñoz, Darío García de Viedma, Laura Pérez-Lago, Belén Rodríguez-Sánchez

**Affiliations:** Clinical Microbiology and Infectious Diseases Department, Hospital General Universitario Gregorio Marañón, Madrid, Spain; Instituto de Investigación Sanitaria Gregorio Marañón, Madrid, Spain; CIBER de Enfermedades Respiratorias (CIBERES CB06/06/0058), Madrid, Spain; Medicine Department, Faculty of Medicine, Universidad Complutense de Madrid, Madrid, Spain

**Keywords:** Fourier-Transform Infrared Spectroscopy, FT-IR, *Klebsiella oxytoca*, *Klebsiella michiganensis*

## Abstract

Healthcare-associated infections (HAIs) are a significant concern worldwide due to their impact on patient safety and healthcare costs. *Klebsiella* spp., particularly *Klebsiella pneumoniae* and *Klebsiella oxytoca*, are frequently implicated in HAIs and often exhibit multidrug resistance mechanisms, posing challenges for infection control. In this study, we evaluated Fourier-transform Infrared (FT-IR) spectroscopy as a rapid method for characterizing a nosocomial outbreak caused by VIM-1-producing *K. oxytoca*.

A total of 47 isolates, including outbreak strains and controls, were collected from Hospital Universitario Gregorio Marañón, Spain and the University Hospital Basel, Switzerland. FT-IR spectroscopy was employed for bacterial typing, offering rapid and accurate results compared to conventional methods like pulsed-field gel electrophoresis (PFGE) and correlating with whole-genome sequencing (WGS) results. The FT-IR spectra analysis revealed distinct clusters corresponding to outbreak strains, suggesting a common origin.

Subsequent WGS analysis identified *Klebsiella michiganensis* as the causative agent of the outbreak, challenging the initial assumption based on FT-IR results. However, both FT-IR and WGS methods showed high concordance, with an Adjusted Rand index (AR) of 0.882 and an Adjusted Wallace coefficient (AW) of 0.937, indicating the reliability of FT-IR in outbreak characterization.

Furthermore, FT-IR spectra visualization highlighted discriminatory features between outbreak and non-outbreak isolates, facilitating rapid screening in case and outbreak is suspected.

In conclusion, FT-IR spectroscopy offers a rapid and cost-effective alternative to traditional typing methods, enabling timely intervention and effective management of nosocomial outbreaks. Its integration with WGS enhances the accuracy of outbreak investigations, demonstrating its utility in clinical microbiology and infection control practices.

## Introduction

Nosocomial infections, also known as healthcare-associated infections (HAIs), are a significant concern within the healthcare system (1). They pose a substantial threat to patient safety, as they can lead to prolonged hospital stays, with increased morbidity and mortality rates. Two of the microorganisms commonly involved in HAIs belong to the *Klebsiella* spp. genus, specifically Kle*bsiella pneumoniae* and *Klebsiella oxytoca* (2). They usually exhibit multidrug resistance mechanisms, such as extended-spectrum β-lactamase or carbapenemase (3). In Spain, the reported rate of carbapenem resistance in *K. pneumoniae* in 2021 was 5.9%, showing an increasing trend (4). For *K. oxytoca* complex, VIM-1 was the most commonly detected carbapenemase (5).

Rapid typing and characterization of bacterial isolates is of high importance for a successful outbreak control and management by detecting transmission routes and environmental reservoirs. Although pulsed-field gel electrophoresis (PFGE) has been conventionally used for bacterial typing, this is a laborious method and requires expertise. In the last years, whole-genome sequencing (WGS) has gained ground as a reference method due to its high discriminatory power and accuracy. However, it is a costly method and requires specific equipment and specialized personnel (6).

Fourier-transform Infrared (FT-IR) spectroscopy, on the other hand, is a rapid and cost-effective method for bacterial typing based on the absorbance patterns of infrared light by different functional groups (7). This methodology has been applied recently for the characterization of outbreaks caused by different bacterial species, offering an accurate and rapid alternative to molecular methods (8–10).

The aim of this study was the evaluation of FT-IR for the characterization of a nosocomial outbreak caused by VIM-1 *K. oxytoca*.

## Materials and methods

### Bacterial strains

Between 2020 and 2022, the Hospital Universitario Gregorio Marañón (HGM; Madrid, Spain) detected an increase in the number of VIM producing *K. oxytoca* isolates sourcing from the children’s ward. A total of 27 isolates identified between 2020 and 2022 as *K. oxytoca* by MALDI-TOF MS and harbouring the VIM-1se resistance mechanism were included in this study. Besides, two control isolates from the 2020-2022 period were also added: one carbapenem susceptible isolate and one isolate harbouring OXA-48 resistance mechanism. Finally, six VIM-1 *K. oxytoca* strains isolated in previous years from adult patients and 12 strains of *K. michiganensis* obtained in University Hospital Basel (Switzerland) (11) were included as controls (Table 1).

**Table 1.**
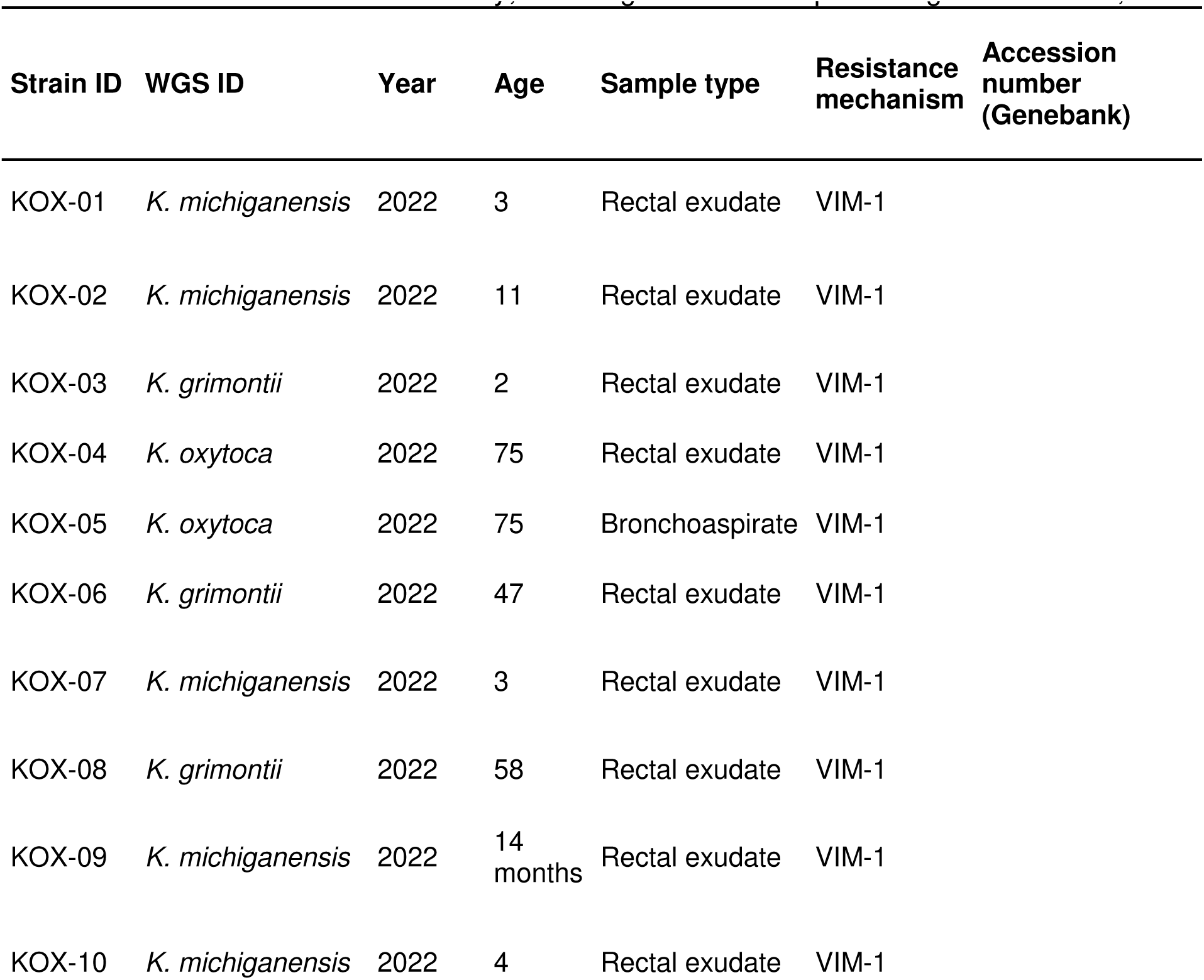
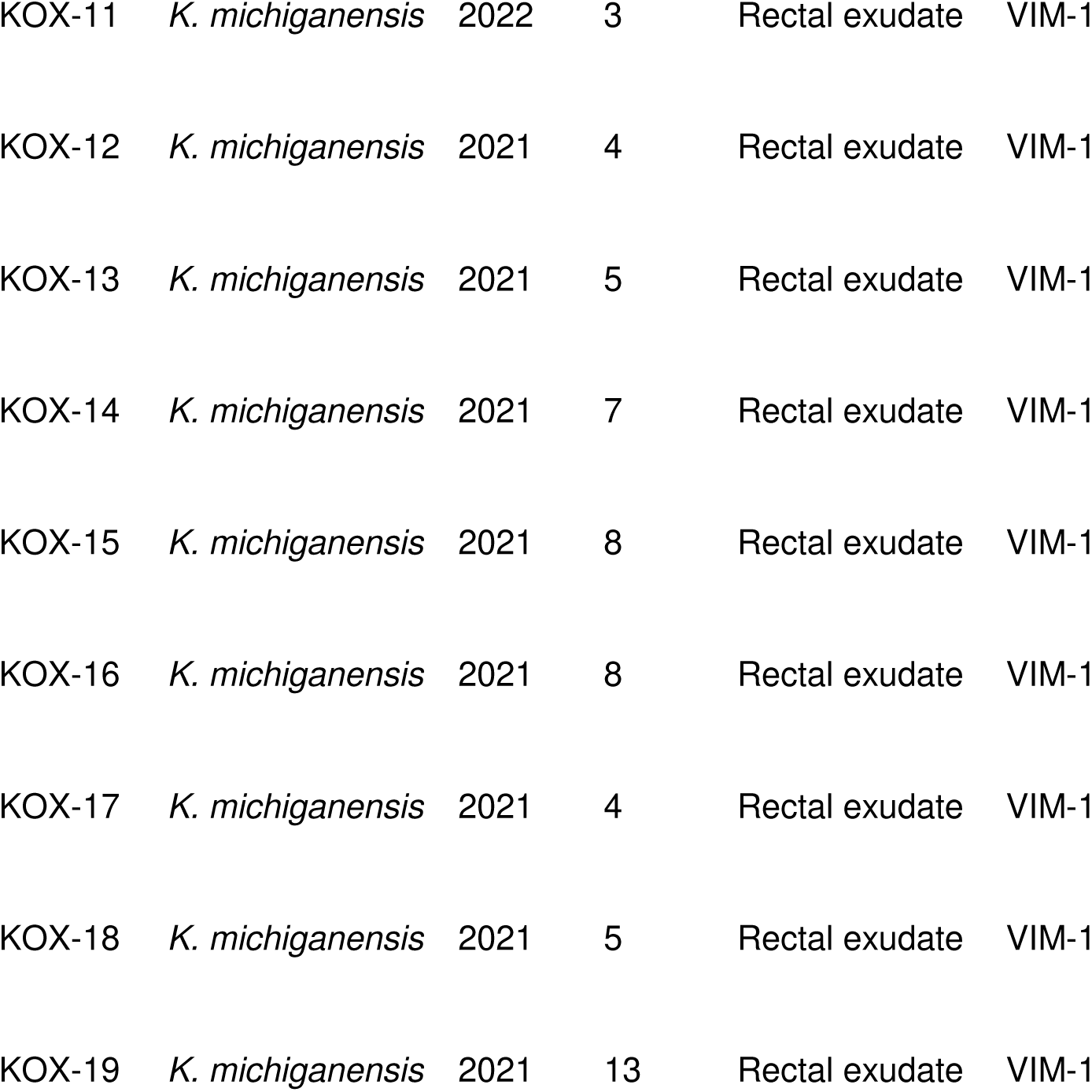
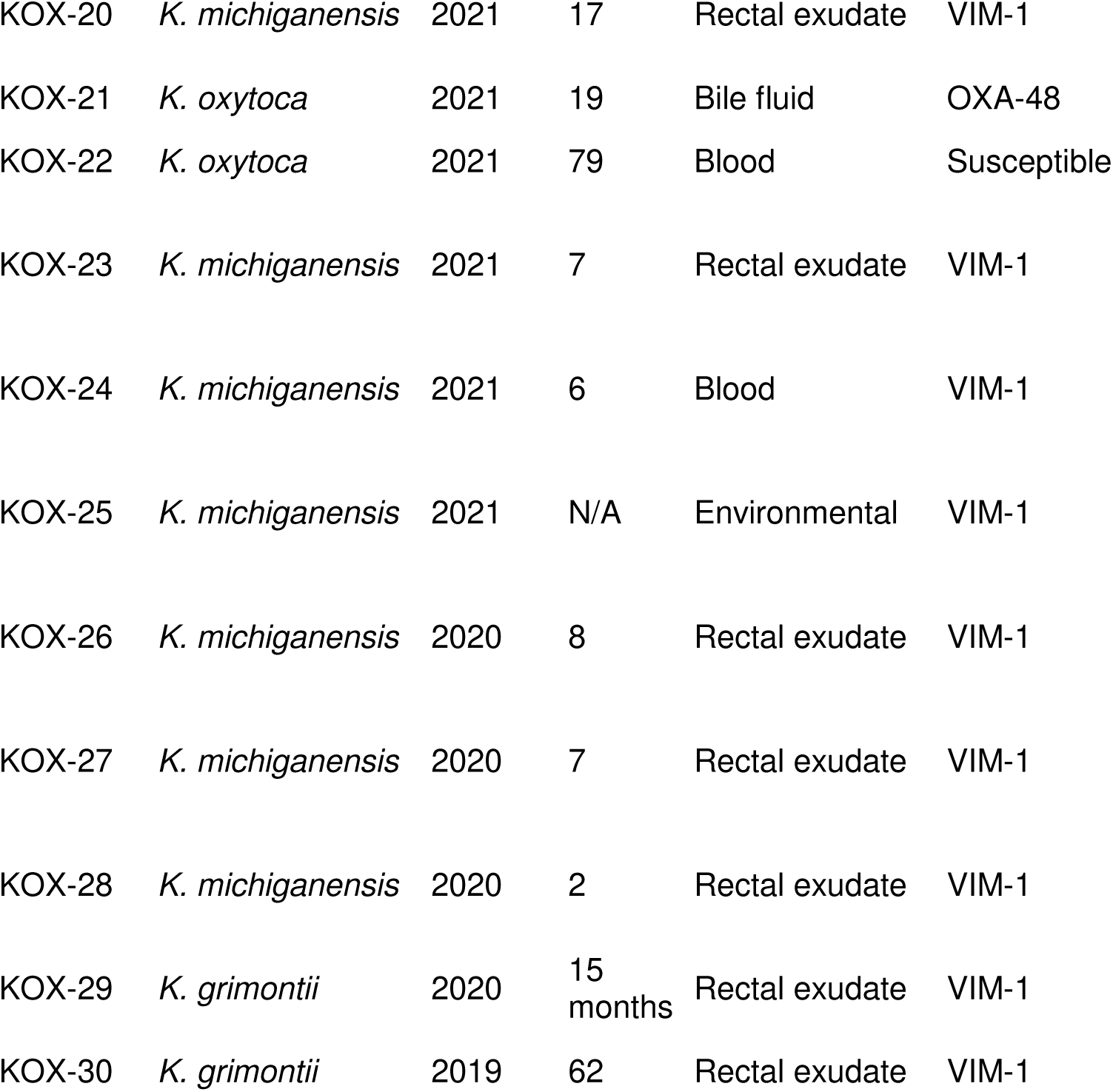
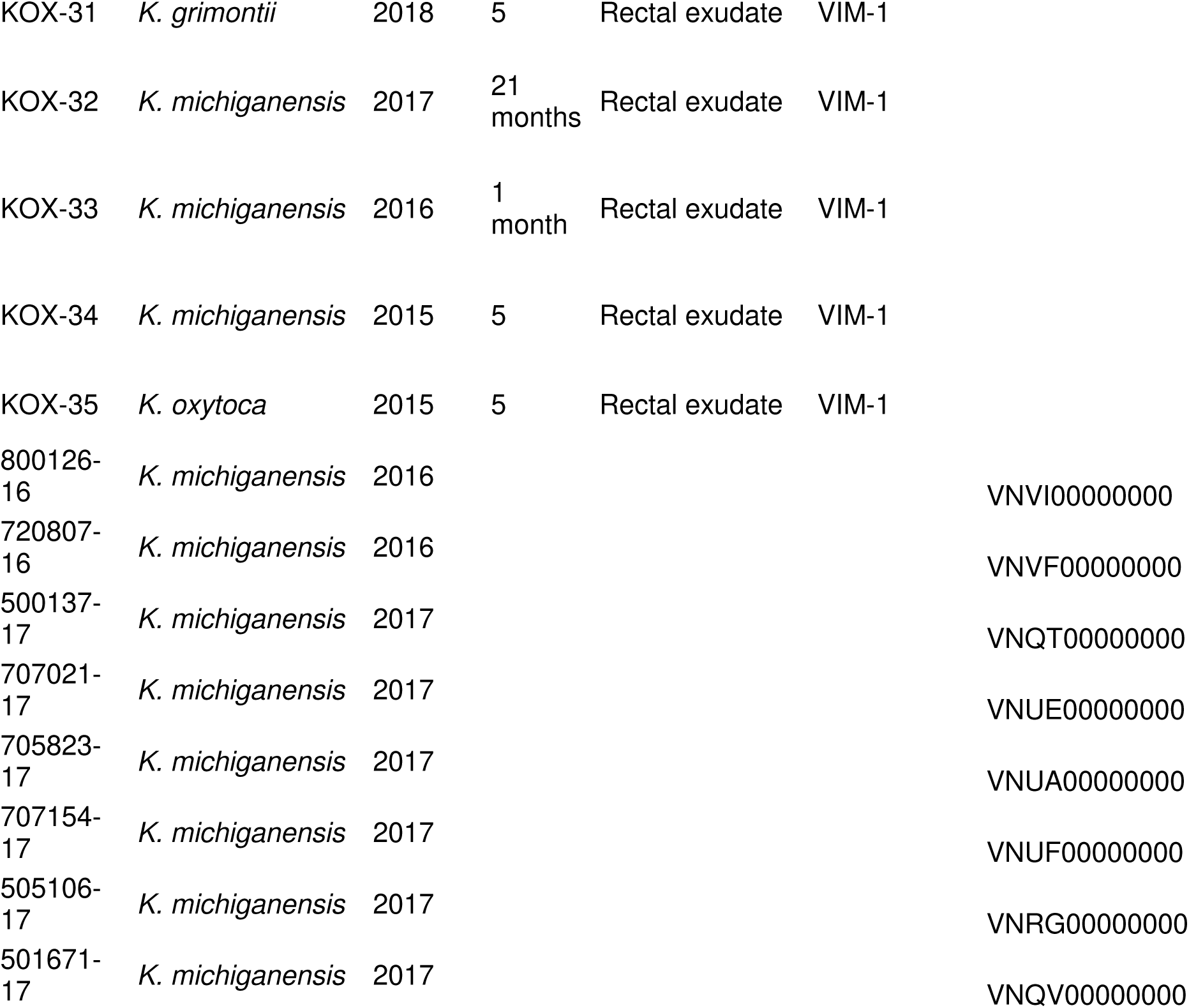
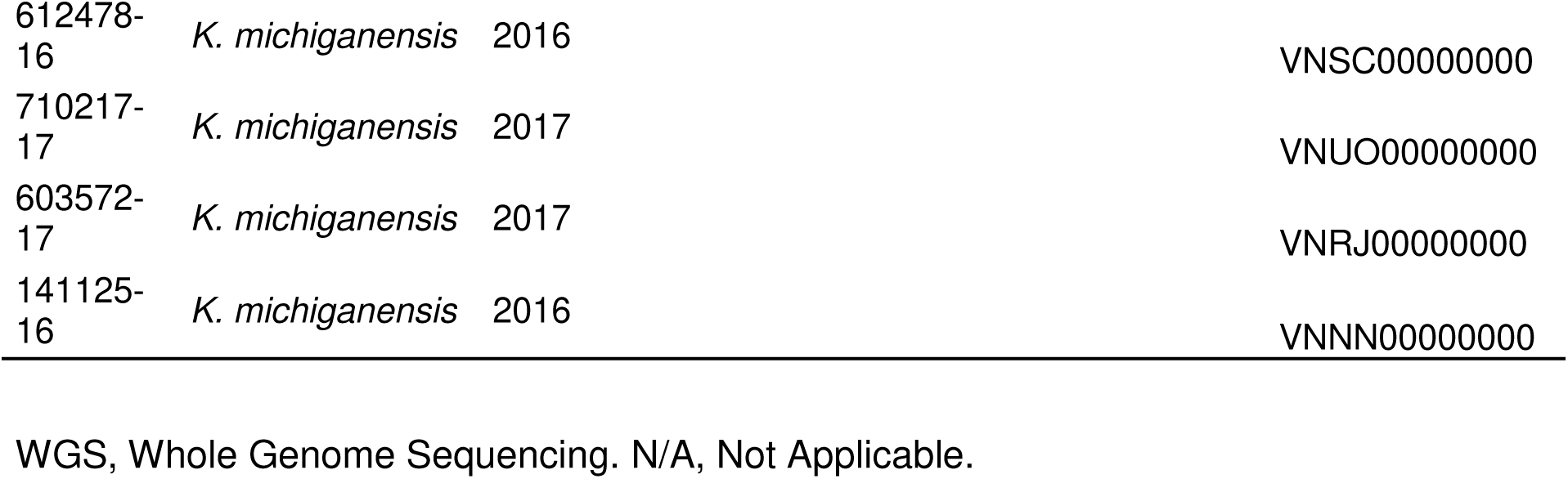
Isolates included in the study, sourcing from the Hospital Gregorio Marañón, Madrid.

### FT-IR spectroscopy

All isolates were recovered from frozen stocks (−80 °C) and cultured on Columbia sheep-blood agar at 37 °C during 24 h. A loop of biomass was resuspended in tubes containing 50 µl of 70% ethanol and metallic rods. The tubes were vortexed for homogenization and 50 µl of HPLC water were added. After vortexing again, 15 µl of the suspension were spotted by triplicate onto the IR Biotyper sample plate (Bruker Daltonics, Bremen, Germany). Samples were analysed in three independent experiments and the spectra that passed all quality controls were further analysed. Using the IR Biotyper software, the mean average spectra for each isolate was calculated. The spectra were processed in the polysaccharide region (1300-800 cm^−1^) applying smoothing by Savitzky-Golay filter (9 points and polynomial degree 3), second derivative and vector normalization. Hierarchical Cluster Analysis (HCA), using Euclidean distance and average linkage, and Principal Component Analysis (PCA) were applied to analyse the distance among spectra. The overall time for sample preparation, analysis and interpretation of the data was around 3 hours. For spectra visualization and representation of plots, we used Clover MS Data Analysis Software (Clover Biosoft, Granada, Spain) and Quasar free software (12).

### Whole-genome sequencing

Nextera XT DNA Sample Preparation Kit (Illumina, San Diego, CA, USA) was used to prepare the libraries for sequencing. Library quality and size distribution were checked on a 2200 TapeStation Bioanalyzer (Agilent Technologies, Santa Clara, US). Libraries were run on the MiSeq system (2×151bp, Illumina), which yielded an average coverage per base of 44.48x (ranging from 8.61 to 87.34x), with 71.80% of the genome having >20x coverage against the reference genome, *K. oxytoca* complex (NZ_CP033844.1).

Sequence analyses were performed using a homemade pipeline deposited in Git-Hub: https://github.com/MG-IiSGM/autosnippy. Briefly, species identification was conducted with Kraken2 v2.1.2 and Mash v2.3. Employing Snippy v4.6.0: mapping was performed with Burrows-Wheeler Alignment (BWA-MEM v0.7.17), and variant calling was performed with FreeBayes v1.3.2, using *K. oxytoca* complex (NZ_CP033844.1) as the reference. Finally, SNPs distances between sequences were calculated using Jaccard similarity and Hamming distance, generating distance matrices and hierarchical dendrograms.

### Correlation between FT-IR and WGS clusters

The clustering consensus obtained by WGS and FT-IR was calculated by the Adjusted Rand index (AR) and Adjusted Wallace coefficient (AW) with 95% CI using the online tool Comparing Partitions (http://www.comparingpartitions.info/). The AR compares the rate of agreement of two methods, and AW considers one of them as the reference method -WGS in this case-. For both indexes, a value of 1 indicates perfect correlation.

## Results

A total of 47 isolates were collected for this study (Table 1). Among the 27 isolates *K. oxytoca*-VIM from the outbreak period, 22 sourced from infant patients suspected to be involved in the outbreak and one was an environmental isolate cultured from a sink in a patients’ rooms; the remaining 4 isolates were obtained from adult patients. All but one of suspected-outbreak isolates were obtained from rectal exudates, implying colonization of patients, and one *K. oxytoca*-VIM was isolated from blood cultures, which implied actual infection.

A dendrogram was built with the FT-IR spectra from the analysed isolates after applying a dimensionality reduction to 8 principal components reaching the 95.2% of variance (Figure 1A). Among the 23 suspected-outbreak isolates, 19 formed a central cluster, including the environmental isolate, whereas the 6 control strains, appeared among the outgroups of the dendrogram. The 4 isolates from adult patients obtained during the outbreak period were also allocated outside the main cluster. According to first epidemiologic information of the isolates, it was considered that the outbreak involved the 17 patients (18 clinical isolates) from the central cluster, in addition to the environmental isolate, which could have been the origin of the outbreak. On the other hand, the outer group with the remaining 4 isolates suggested that these patients were not related to the outbreak although their *K. oxytoca* isolates were detected simultaneously to the outbreak. The central cluster was separated from the other strains by a distance of 0.32-0.42, as shown in the distance matrix (Figure 1B). Through the PCA visualization (Figure 1C), a diffuse cluster was observed on the left, that corresponded to the strains of the central cluster in HCA. Most of the not-related strains appeared along the right region of the PCA.

**Figure 1.**
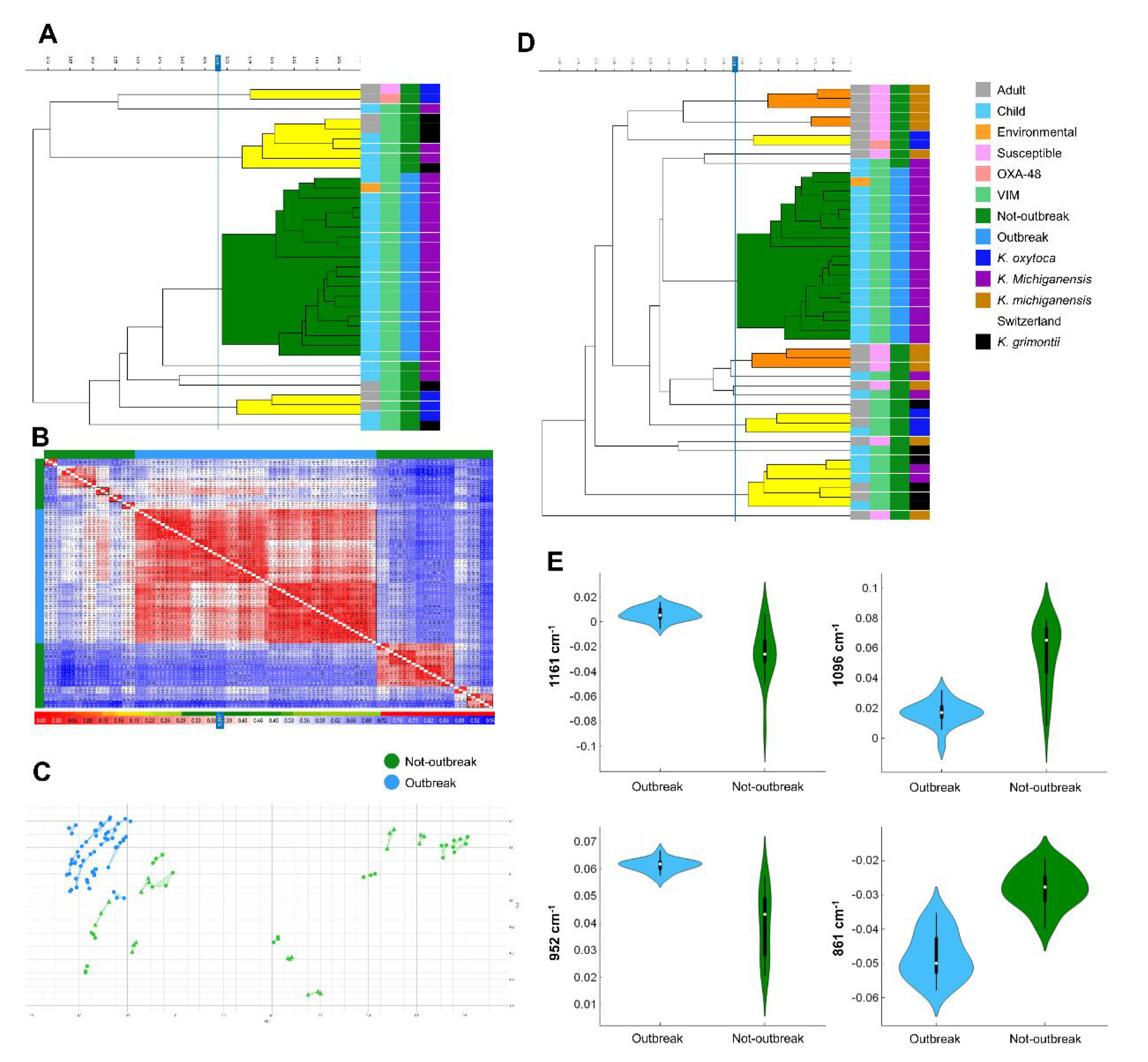
Analysis of Fourier-Transform Infrared spectra of the region 1300-800 cm^−1^, using IR Biotyper and Quasar software. **A**, Dendrogram with the 35 strains obtained in Hospital Gregorio Marañón, according to Whole-Genome Sequencing (WGS), outbreak isolates are those of the green cluster. **B**, Distance matrix of 35 strains, low distance spectra are red-coloured, and high distance spectra are blue-coloured. The central red cluster corresponded to the outbreak strains, indicating a low distance between them. **C**, Principal Component Analysis of the 35 isolates. A cluster was observed at the left of the image; after WGS it was confirmed which of them corresponded to the outbreak. **D**, Dendrogram with 12 *K. michiganensis* strains from Switzerland as outgroups added to the initial analysis. **E**, Wavelength regions of the spectra with higher difference in absorbance values between outbreak and not-outbreak isolates: 1161 cm^−1^, 1096 cm^−1^, 952 cm^−1^ and 861 cm^−1^.

After performing of WGS, different species from the *K. oxytoca* complex were identified: *K. oxytoca*, *Klebsiella michiganensis* and *Klebsiella grimontii*. The predominant species was *K. michiganensis*, which was only isolated from children, and was identified as the actual cause of the outbreak (Table 1). Comparative analysis of 21 *K. michiganensis* strains from the study period revealed that 19 had pairwise distances of 0-16 SNPs, confirming their outbreak nature. These 19 isolates corresponded to those grouped in the central cluster by FT-IR. The remaining 2 *K. michiganensis* strains exhibited >10,000 SNP differences, excluding them from the outbreak. Comparing WGS with FT-IR clustering, we obtained an AR index of 0.882 (95% CI, 0.739-1.000) and an AW value of 0.937 (95% CI, 0.897-0.978). To analyse the robustness of the previous classification, 12 *K. michiganensis* strains sourcing from the University Hospital Basel (Switzerland) were included as an outgroup (11), analysed by FT-IR spectroscopy and compared with the spectra from the initial clustering described above. The dendrogram obtained is shown in Figure 1D. The spectra from the outbreak samples remained a distinct cluster as in the first analysis, whereas the rest of the strains were redistributed.

The spectra were further visualized to find differences between the outbreak strains and the not-related isolates. As can be observed in Figure 2, the spectra from the outbreak strains differed only slightly among them, whereas they showed differences when compared with strains belonging to *K. grimontii* (Figure 2A) and *K. oxytoca* (Figure 2B) species. In comparison with other strains from the same species, outbreak strains also showed visual differences with other *K. michiganensis* isolated during the same period (Figure 2C) and with *K. michiganensis* strains from Switzerland (Figure 2D). Overall, the most discriminative regions of the spectra were located at wavelengths of 1161, 1096, 952 and 861 cm^−1^ (Figure 1E).

**Figure 2.**
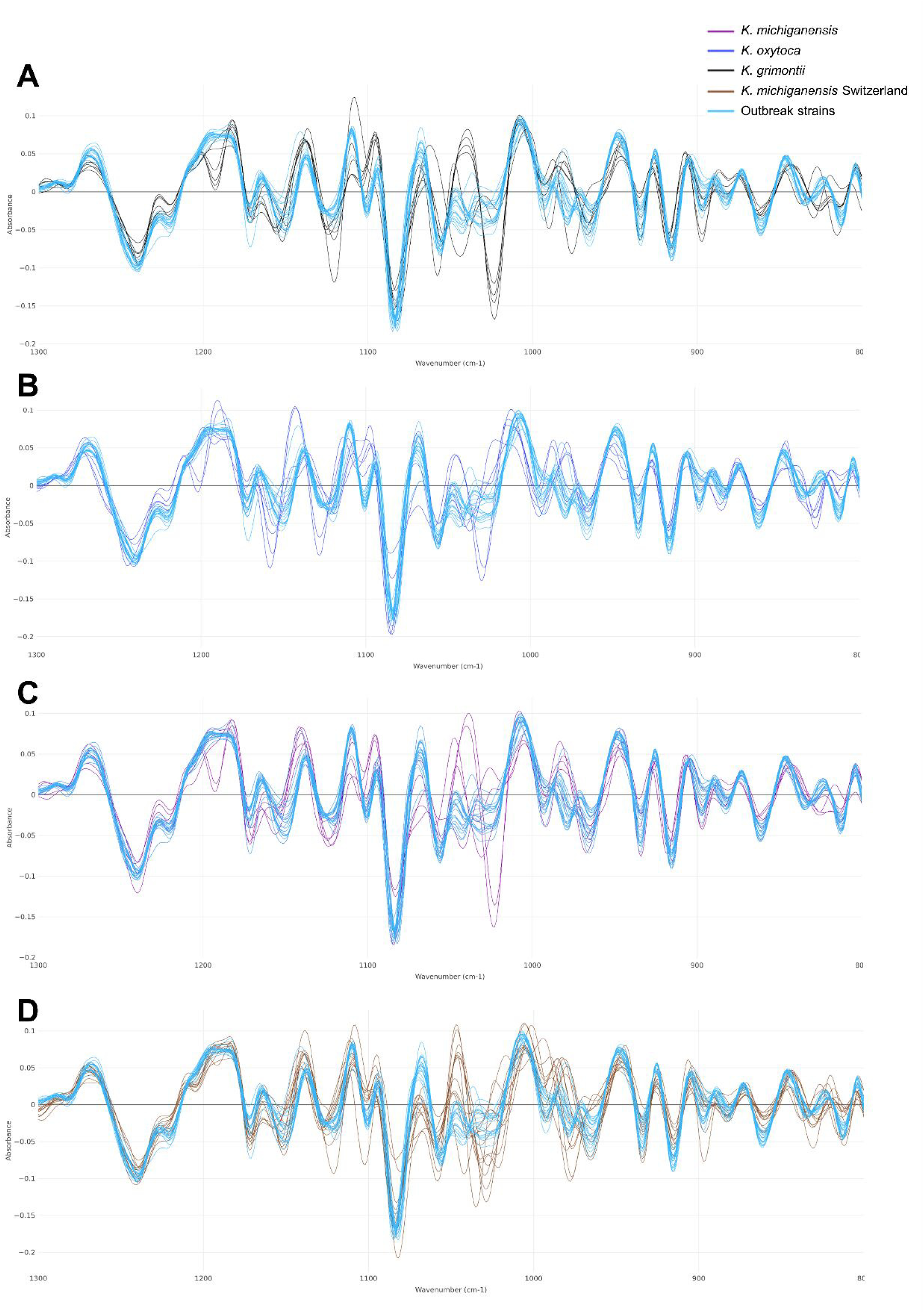
Spectra region of 1300-800 cm^−1^ corresponding to the polysaccharides. The strains involved in the outbreak (light blue) are compared with other groups of strains of the study. Spectra visualization was performed by Clover MS Data Analysis Software: **A,** *K. grimontii* spectra (black); **B**, *K. oxytoca* spectra (dark blue); **C**, *K. michiganensis* spectra (violet); **D**, spectra of *K. michiganensis* obtained from Switzerland (brown).

## Discussion

*K. oxytoca* has been reported to be involved in several HAIs (1). Moreover, these outbreaks were often related with neonatal intensive care units and can imply carbapenem-resistant strains (13, 14). In the present study, an increase of VIM-producing *K. oxytoca* was detected during the 2020-2022 period in rectal exudates from patients in the children’s ward of a tertiary hospital. Although most of the patients were colonized, one of them developed bacteraemia by this microorganism.

The first analysis, performed by FT-IR spectroscopy, showed that 19 of the 23 strains suspected to be involved in the outbreak formed a central cluster in the dendrogram, which also included one environmental isolate. This result suggested the environmental origin of the outbreak. Four other isolates sourcing from infant patients in the same period, and suspected to be also involved in the outbreak, were distributed in other regions of the FT-IR dendrogram, along with the negative controls and the isolates from adult patients. Therefore, these 4 patients could be considered as unrelated to the outbreak.

Although the microorganism most frequently analyzed by FT-IR spectroscopy is *K. pneumoniae*, the cut-off used for strain typing differed between studies. Whereas Martak et al. defined a wide cut-off between 0.181-0.864 (15), other authors decreased this range to 0.196-0.439 (10, 16), which is in consonant with our study, although in this case the outbreak was caused by *K. oxytoca*, and the cut-off values are known to vary between species. In fact, to our knowledge, only one previous study applied FT-IR spectroscopy for typing of *K. oxytoca* in a context of liquid hand soap contamination, with successful classification but no comparable cut-off (17).

By the application of WGS, the first finding was that the outbreak was not caused by *K. oxytoca*, but *K. michiganensis*, a species within the *K. oxytoca* complex (18). Moreover, among the isolates not related to the outbreak, other members of the same complex -*K. michiganensis* and *K. grimontii*-were identified. In fact, the identification performed by MALDI-TOF MS only reached complex-level for *K. pneumoniae* and *K. oxytoca* complexes, although some efforts have been made for a more accurate species-level identification (19). Therefore, the real incidence of *K. michiganensis* could be underestimated. WGS confirmed the presence of VIM-1 carbapenemase on an IncFIB(pQIL) plasmid in all *K. michiganensis* outbreak isolates, a mechanisms recently described (20). The short genomic distances found among the 19 isolates included in the outbreak by FT-IR spectroscopy confirmed the linkage between these strains, while excluding the linkage of 2 strains despite their temporal coincidence with the paediatric patients involved in the outbreak.

In conclusion, FT-IR spectroscopy showed high correlation with WGS for the characterization of this outbreak. The analysis by FT-IR spectroscopy implies a simple procedure, is rapidly performed and generates accurate results. Thus, the proposed workflow could be used as a first screening for potential outbreaks and for the selection of interesting isolates for a better characterization by WGS techniques.

## Acknowledgements

The authors are grateful to Dr. Adrian Egli and Dr. Aline Cuénod, currently belonging to the Institute of Medical Microbiology, University of Zurich, Switzerland, and the McGill University, Montréal, Canada, respectively, for providing *K. michiganensis* isolates from the University Hospital Basel.

This work is partially supported by grant PI23/01288 funded by the Health Research Fund of the Carlos III Health Institute (ISCIII, Madrid, Spain) partially financed by the European Regional Development Fund (FEDER) ‘A way of making Europe’. The funders had no role in the study design, data collection, analysis, decision to publish, or preparation/content of the manuscript. DRT (Sara Borrell CD22-00014) and LPL (Miguel Servet CPII20/0001) are funded by ISCIII. MBS is the recipient of the Intramural predoctoral contract 2022 from the Health Research Center of the Hospital Gregorio Marañón -IISGM-.

